# Conservation and divergence of transcriptional heterogeneity in the cardiac conduction system

**DOI:** 10.1101/2025.09.23.678069

**Authors:** Marwan Bakr, Saif Dababneh, Glen F. Tibbits, Yena Oh, Kyoung-Han Kim

## Abstract

The cardiac conduction system (CCS) consists of specialized cardiomyocytes that initiate and propagate electrical activity through the heart. While the transcriptional programs underlying CCS development and function have been studied within individual species, how these programs compare across species and developmental stages remains unclear. Here, we present a comprehensive cross-species and cross-stage analysis of the CCS transcriptome using single-cell/single-nucleus RNA sequencing and spatial transcriptomic datasets from human, mouse, rat, zebrafish, and medaka hearts. We identify shared and species-or stage-specific gene expression patterns, spanning CCS-wide, zonal-, and component-level features, as well as conserved gene regulatory networks across the species. Many conserved genes are associated with human CCS function and related disorders, highlighting their translational relevance for conduction disease. This work refines the molecular characterization of CCS cell types across vertebrates and provides a resource for advancing our understanding of CCS development, function, and pathology.

## INTRODUCTION

The cardiac conduction system (CCS) is composed of specialized cardiomyocyte populations responsible for initiating and propagating electrical signals throughout the heart. The development and distinct function of each CCS component rely on specialized gene expression profiles. For example, *Hcn4* is highly expressed in the sinoatrial node (SAN) and is responsible for its automaticity^1, 2^, while Cx40, a gap junction protein encoded by *Gja5*, is enriched in the ventricular conduction system (VCS), enabling its rapid conduction^3^. These heterogeneous gene programs within the CCS are established and maintained by networks of transcription factors (TFs), such as Shox2^4^, Isl1^5^ and Tbx18^6^ in SAN; Tbx3^7^ in SAN, atrioventricular node (AVN) and the proximal VCS (*His* bundle and bundle branches); and Nkx2-5^8^, Etv1^9^ and Irx3^10, 11^ in the VCS. Importantly, disruptions of these key TFs often lead to electrophysiological abnormalities in mice, comparable to arrhythmias and conduction abnormalities present in humans. For example, the absence of *Shox2* or *Isl1* results in bradycardia, while deficiencies in *Nkx2-5* or *Irx3* cause prolonged QRS duration and bundle branch block in mice^12^. Genetic variants in these TFs and their downstream target genes have also been linked to human conduction disorders^12, 13^, indicating a general conservation of CCS gene programs.

Several model organisms are widely used to study the CCS, as overall patterns of electrical impulse generation, propagation, and structural organization are comparable across species. However, species-specific physiological differences, such as heart rate, size, and anatomy, can complicate the direct translation of findings from model organisms to human physiology and disease. In addition, disparities exist between the developing and mature hearts, and our knowledge of CCS maturation at the molecular level remains limited. Our recent work has demonstrated that despite global transcriptomic similarities between the embryonic and early postnatal mouse CCS, there are shifts in molecular programs during this transition^14^. Moreover, the presence of proliferating cells in the postnatal mouse CCS^14^, and the hypoplastic phenotypes observed in the postnatal, but not embryonic, CCS lacking key TFs (e.g., *Irx3*, *Tbx3*, and *Nkx2.5*) suggest continued maturation of the CCS after birth.

Advances in single-cell and single-nucleus RNA sequencing (sc/snRNA-seq) and spatial transcriptomics have begun to uncover the transcriptomic profiles of the CCS in various species^14–22^. However, most studies have focused on individual CCS components, primarily the SAN, rather than CCS-wide heterogeneity, and have lacked systematic cross-species analyses. Furthermore, CCS populations are often not annotated in broader cardiac sequencing studies.

Together, these gaps highlight the need for systematic, cross-species, and cross-stage analyses of the transcriptional programs to gain a deeper understanding of both conserved and divergent features of CCS development and function. This knowledge is critical for several translational applications, including elucidating the pathological mechanisms of conduction diseases using animal models, developing therapeutic strategies targeting specific CCS regions, and reprogramming stem cells into CCS-like cells, such as the development of biological pacemakers.

In this study, we conduct a comparative analysis to enhance our understanding of the conservation and divergence of CCS transcriptional heterogeneity across species and development. We identify evolutionarily conserved regional CCS markers, as well as those specific to a particular species and/or developmental stages. Furthermore, we investigate the gene regulatory networks underlying the heterogeneous expression of these conserved gene programs and assess their translational relevance by examining associations between genetic variants and conduction disorders.

## RESULTS

### Identification of CCS cell populations in humans, mice, rats, and zebrafish

To compare the transcriptomic profiles of CCS components across species and developmental stages, we utilized nine single-cell and single-nucleus transcriptomic datasets from humans^18–20^, mice^14, 21, 22^, rats, zebrafish^15–17^, and medaka^15^, at developing and mature stages (**Fig. 1a**). We also used spatial transcriptomics data, including MERFISH imaging of human fetal whole heart sections^19^ (12 – 13 post-conceptional weeks; pcw) and Visium slides of various cardiac regions from adult human donors^20^. Cell annotations were either obtained directly from the original studies, assigned using the methods described therein, or labelled independently where necessary (i.e., fetal human in **Supplementary** Fig. 1, adult rat in **Fig. 4**, and adult zebrafish hearts in **Supplementary** Fig. 2). A summary of all datasets and annotation approaches is provided in **Supplementary Table 1**.

**Figure 1.**
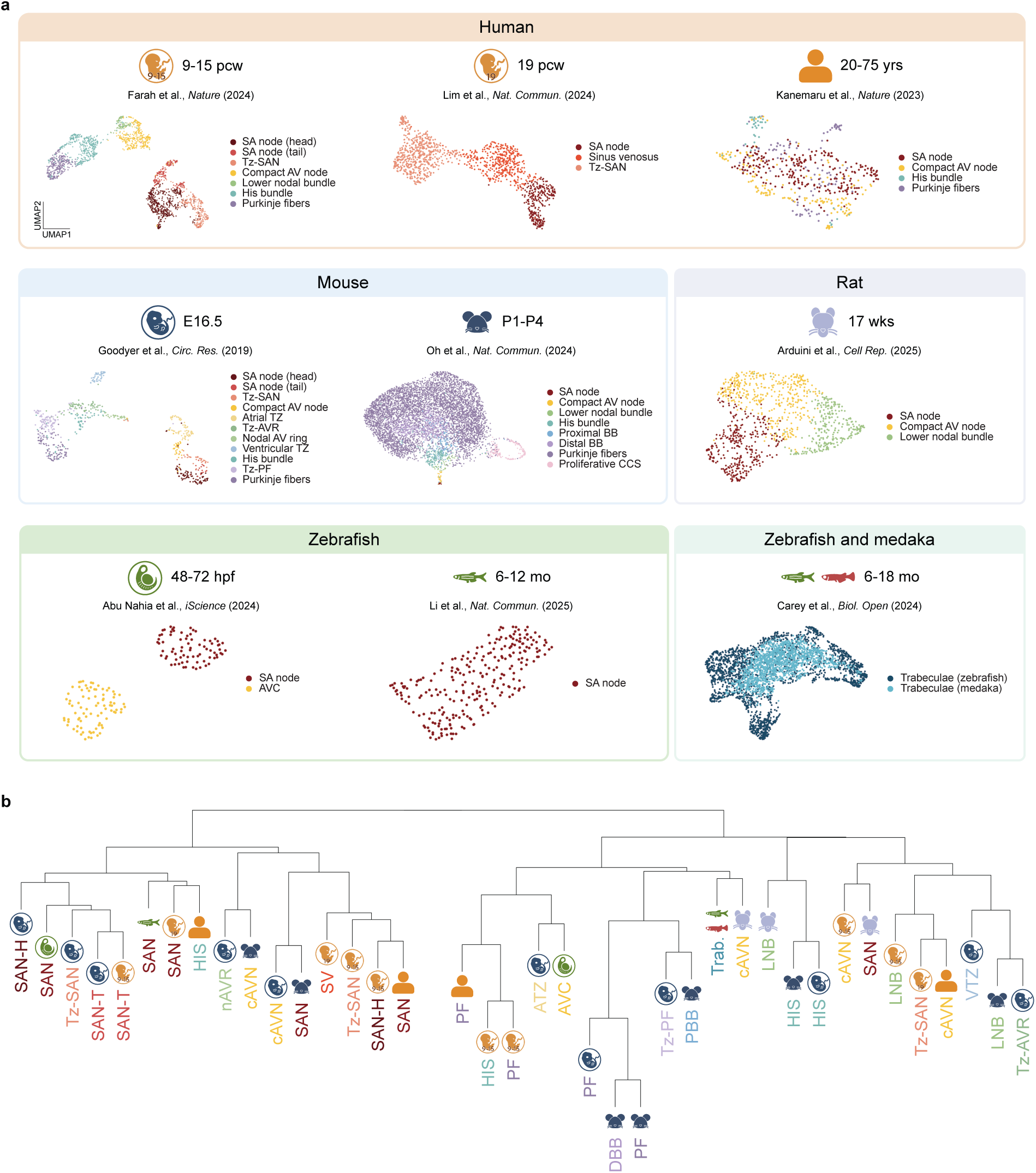
Single-cell/nucleus transcriptomic profiles of the cardiac conduction system (CCS) across vertebrate species. **a**, Uniform manifold approximation and projection (UMAP) embeddings of CCS cells computationally isolated from single-cell/nucleus RNA sequencing datasets (see Supplementary Table 1 for details). **b**, Hierarchical clustering of CCS components based on Pearson correlation of average expression profiles. SAN, sinoatrial node; SAN-H, sinoatrial node head; SAN-T, sinoatrial node tail; Tz-SAN, transitional sinoatrial node; SV, sinus venosus; cAVN, compact atrioventricular node; AVC, atrioventricular canal; nAVR, nodal atrioventricular ring; Tz-AVR, transitional atrioventricular ring; LNB, lower nodal bundle; ATZ, atrial transitional zone; VTZ, ventricular transitional zone; HIS, *His* bundle; PBB, proximal bundle branch; DBB, distal bundle branch; PF, *Purkinje* fibers; Tz-PF, transitional *Purkinje* fibers; Trab, trabeculae; pcw, post-conception weeks; yrs, years; wks, weeks; mo, months; E16.5, embryonic day 16.5; P1-P4, postnatal days 1-4; hpf, hours post-fertilization.

In datasets we annotated independently, CCS components were defined based on well-established marker genes, yielding comparable profiles across all datasets. The SAN was marked by *SHOX2*^4^, *VSNL1*^14, 17, 23^, *TBX3*^7^, and *HCN4*^1, 2^, with mutually exclusive expression of *TBX18* in the head and *NKX2-5* in the tail^6, 24, 25^. The compact atrioventricular node (cAVN) was defined by *TBX3*, *RSPO3*, *BMP2*, and *CACNA1D*^14^, along with the atrial marker *MYL7*. We defined the lower nodal bundle (LNB) based on the expression of AVN genes (e.g., *TBX3*, *BMP2*, *CACNA1D*) as well as ventricular chamber genes (*MYH7*, *MYL2*)^14^. The *His* bundle (HIS) was identified by the expression of proximal VCS markers, such as *TBX3*^7^ and *ROBO1*^26, 27^, while *Purkinje* fibres (PF) were identified by the expression of distal VCS markers, including *IRX3*^10, 11, 28^, *GJA5*^3^, and *SEMA3A*^29^. Transitional cells were defined by hybrid expression of CCS-specific regional markers as well as surrounding contractile cardiomyocyte markers. To facilitate broader access and exploration of CCS-specific transcriptomic data, we have developed an interactive web application (https://ccsatlas.com) that allows users to query gene expression in all datasets included in this study. To assess similarities between CCS components across species and developmental stages, we performed unsupervised hierarchical clustering of CCS populations from all datasets based on gene expression (**Fig. 1b**). Overall, CCS components clustered primarily based on their anatomical regions, indicating significant transcriptomic similarities between corresponding CCS components across species and developmental stages. The clustering distinguished two broad groups: one comprising the SAN along with some cAVN populations and the other encompassing VCS populations, which were further subdivided into proximal (e.g., LNB, HIS) and distal (e.g., PF) VCS components, though with some exceptions.

### Zone-specific CCS transcripts conserved across humans and mice

We first performed cross-species comparisons between humans and mice, as datasets from these two species contain all major CCS components (SAN, cAVN, HIS, and PF) across both developmental and mature stages. Pearson correlation analyses of gene expression profiles showed strong concordance between corresponding CCS components in developing human (9 – 15 pcw) and mouse (embryonic day 16.5; E16.5) hearts, as well as between adult (20 – 75 years) and postnatal (postnatal days 1 – 4; P1 – P4) stages (**Supplementary** Fig. 3). Using these datasets, we identified shared molecular profiles of the global CCS as well as two broad functional zones of the CCS, the nodal components (SAN and cAVN) and the impulse-propagating VCS (HIS and PF).

#### Whole cardiac conduction system

While no universally conserved pan-CCS marker was observed across all four datasets, *ID3*, a gene not previously linked to the CCS, was identified as a pan-CCS marker in developing and adult humans, as well as postnatal mice (**Fig. 2a**). Additionally, the developing human and mouse CCS globally expressed *UNC5B* (a netrin receptor), and *IGFBP5*, which has been shown to be broadly expressed across the developing mouse CCS^22^. Interestingly, *IGFBP5* exhibited higher expression in atrial cardiomyocytes in the postnatal mouse heart but showed limited expression in the adult human CCS.

**Figure 2.**
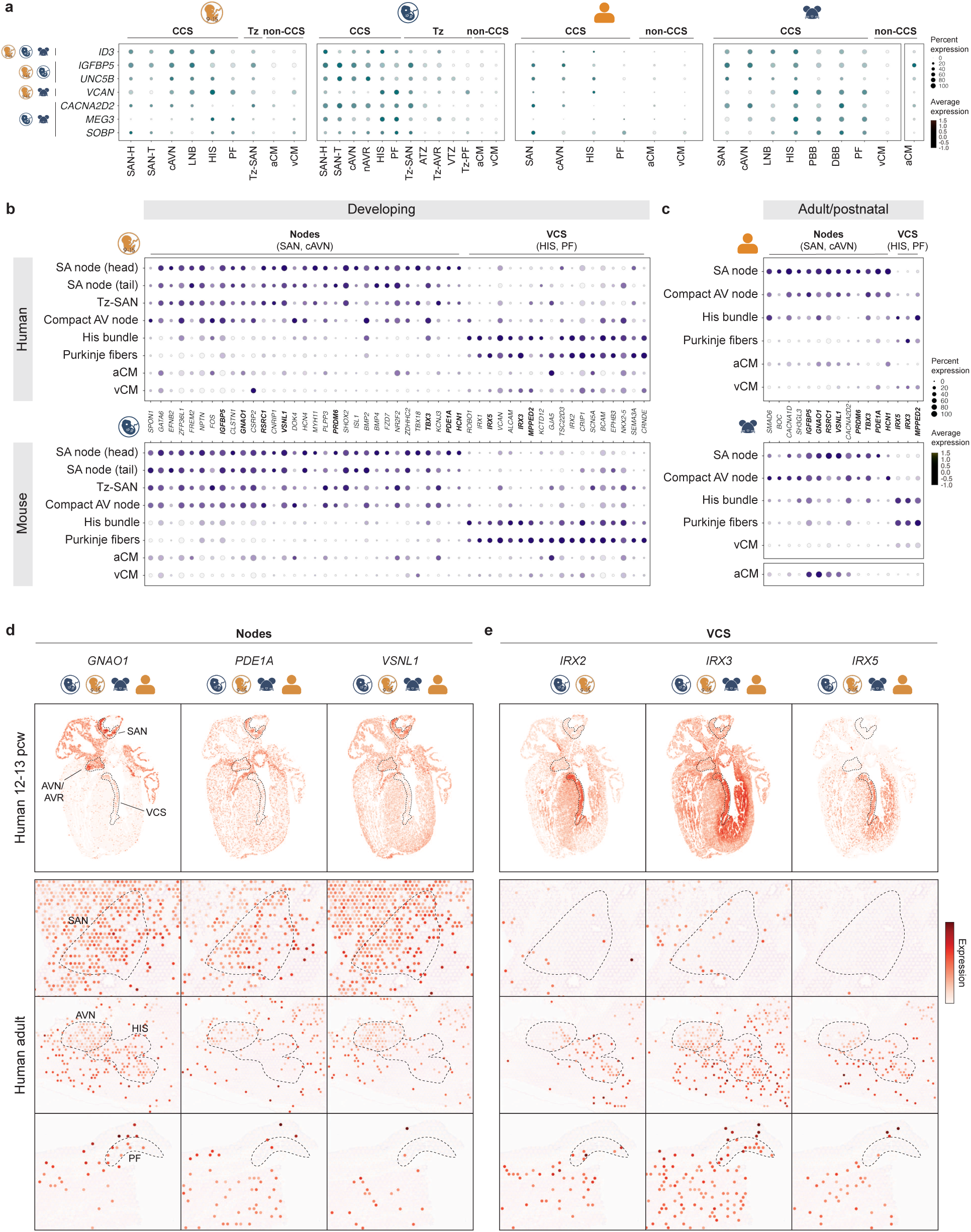
Conserved zonated molecular profiles in the human and mouse CCS. **a**, Expression of species– and stage-shared pan-CCS markers. Mouse postnatal atrial cardiomyocyte data were obtained from a separate dataset (GSE193346). **b-c**, Expression of conserved markers with zonated enrichment in the nodes (SA node and compact AV node) or ventricular conduction system (VCS; *His* bundle and *Purkinje* fibers) in the developing CCS (**b**) and adult/postnatal CCS (**c**). Genes conserved across both developing and adult/postnatal stages are shown in bold. **d-e**, Spatial expression patterns of representative nodal (**d**) and VCS markers (**e**) in a human fetal heart section (12-13 pcw, MERFISH) and from adult heart sections (Visium). aCM, atrial cardiomyocytes; vCM, ventricular cardiomyocytes.

#### Nodes

Genes conserved in the nodal regions between humans and mice at both developing and adult/postnatal stages (**Fig. 2b-c**) included *TBX3*, a TF critical for nodal development^7^; *HCN1*, encoding an ion channel involved in pacemaker activity^30^; *PDE1A*, known to regulate pacemaker activity in rabbits^31^; and *RSRC1*, located adjacent to *SHOX2* in the same genomic locus. Additional genes previously identified and validated in mouse nodes, such as *IGFBP5*^22^, *VSNL1*^14, 23^, *GNAO1*^14, 32^, and *PRDM6*^14^, were also conserved in human nodes. Spatial visualization further supported the expression of *GNAO1*, *PDE1A,* and *VSNL1* in both the developing and adult human nodes (**Fig. 2d**).

We also found multiple genes conserved between human and mouse nodes, specifically during development (**Fig. 2b**). For instance, *GATA6*, which regulates the development of both the SAN and AVN, and harbours variants associated with arrhythmias in humans^33, 34^; *FZD7*, a receptor for Wnt ligands, which have been implicated in SAN development and function^35, 36^; and *BMP2*, known to be expressed in the mouse atrioventricular canal^37^ and cAVN^14^, were enriched in both species. Additional conserved genes, not previously linked to nodal development, included *SPON1*, *CSRP2*, *EFNB2*, *FREM2*, *PLPP3*, and *CNRIP1*. Despite this overall conservation, some genes displayed differences in species-specific expression patterns. For example, *SPON1* and *CSRP2* were strongly expressed in both human and mouse cAVN but displayed weaker expression in the human SAN compared to the mouse SAN.

In the adult/postnatal nodes, conserved genes included *SMAD6*, an inhibitor of BMP signaling; *BOC* and *SH3GL3*, which currently have no known roles in the CCS; as well as *CACNA2D2*^38^ and *CACNA1D,* voltage-gated calcium channel subunits^38, 39^ (**Fig. 2c**). Interestingly, some of these genes exhibited divergent expression patterns outside the nodal regions. For example, *SMAD6* and *CACNA1D*, which were highly specific to mouse nodes, were also expressed in the human HIS. Conversely, *SH3GL3* and *CACNA2D2*, enriched in the human nodes, were expressed in the mouse HIS.

#### Ventricular conduction system

In the VCS, only a few genes were conserved between humans and mice at both developing and adult/postnatal stages (**Fig. 2b-c**). These included *Iroquois* homeobox transcription factors, *IRX3*, which is essential for VCS development and function^10, 11, 40^, and *IRX5*, known to regulate the transmural ventricular gradients of repolarization and contractility in mice^41–43^. In addition to their expression in the VCS, spatial visualization showed broader endomyocardial-enriched expression patterns for both *IRX3* and *IRX5* (**Fig. 2e**). Another conserved gene, *MPPED2*, recently identified as a postnatal mouse VCS marker^14^, was expressed in both developing and adult human VCS. Notably, while *Mpped2* showed stronger expression in the mouse PF^14^, it was most prominently expressed in the HIS at both fetal and adult stages in humans.

In the developing human and mouse VCS, *NKX2-5*, *GJA5*, and *SCN5A* were conserved (**Fig. 2b**), all of which have established roles in VCS development or function^44–47^. Additionally, we observed conserved expression of *IRX1* and *IRX2* (**Fig. 2e**); *CRNDE*, a long non-coding RNA residing in the same genomic locus as *IRX3* and *IRX5*; and *TSC22D3*. However, the specific roles of these genes in the VCS remain to be determined.

### CCS component-specific transcripts conserved across humans and mice

We next identified molecular markers that are specific to each CCS component (SAN, cAVN, HIS, and PF) in human and mouse hearts (**Fig. 3** and **Supplementary Table 2**). Overall, we recognized three groups of marker genes: (i) those conserved across both species and developmental stages, (ii) those conserved across species at comparable developmental stages, and (iii) those conserved across developmental stages within each species.

**Figure 3.**
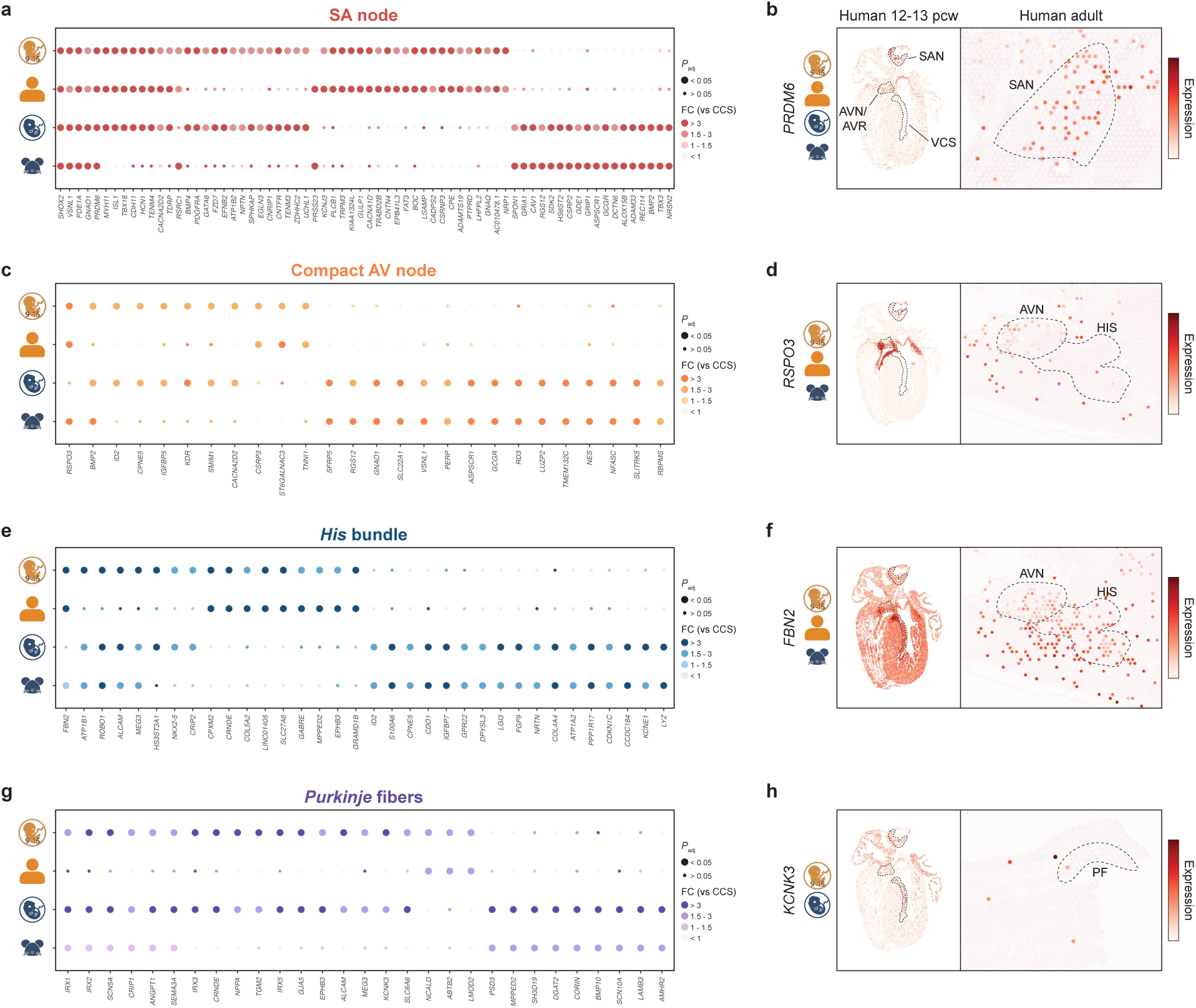
Species– and stage-shared molecular profiles of individual CCS components in humans and mice. Specificity of species– and stage-shared markers for the SA node (**a**), compact AV node (**c**), *His* bundle (**e**), and *Purkinje* fibers (**g**). Dot size represents the significance of differential expression compared to non-CCS cardiomyocytes and to other CCS regions (Bonferroni-adjusted *P*); genes not significant in either comparison are labeled as non-significant. Colour intensity represents the absolute fold change relative to other CCS regions. Spatial visualization of representative marker genes for the SA node (**b**), compact AV node (**d**), *His* bundle (**f**), and *Purkinje* fibers (**h**), in a section of human fetal heart (12-13 pcw, MERFISH) and adult human heart (Visium).

**Figure 4.**
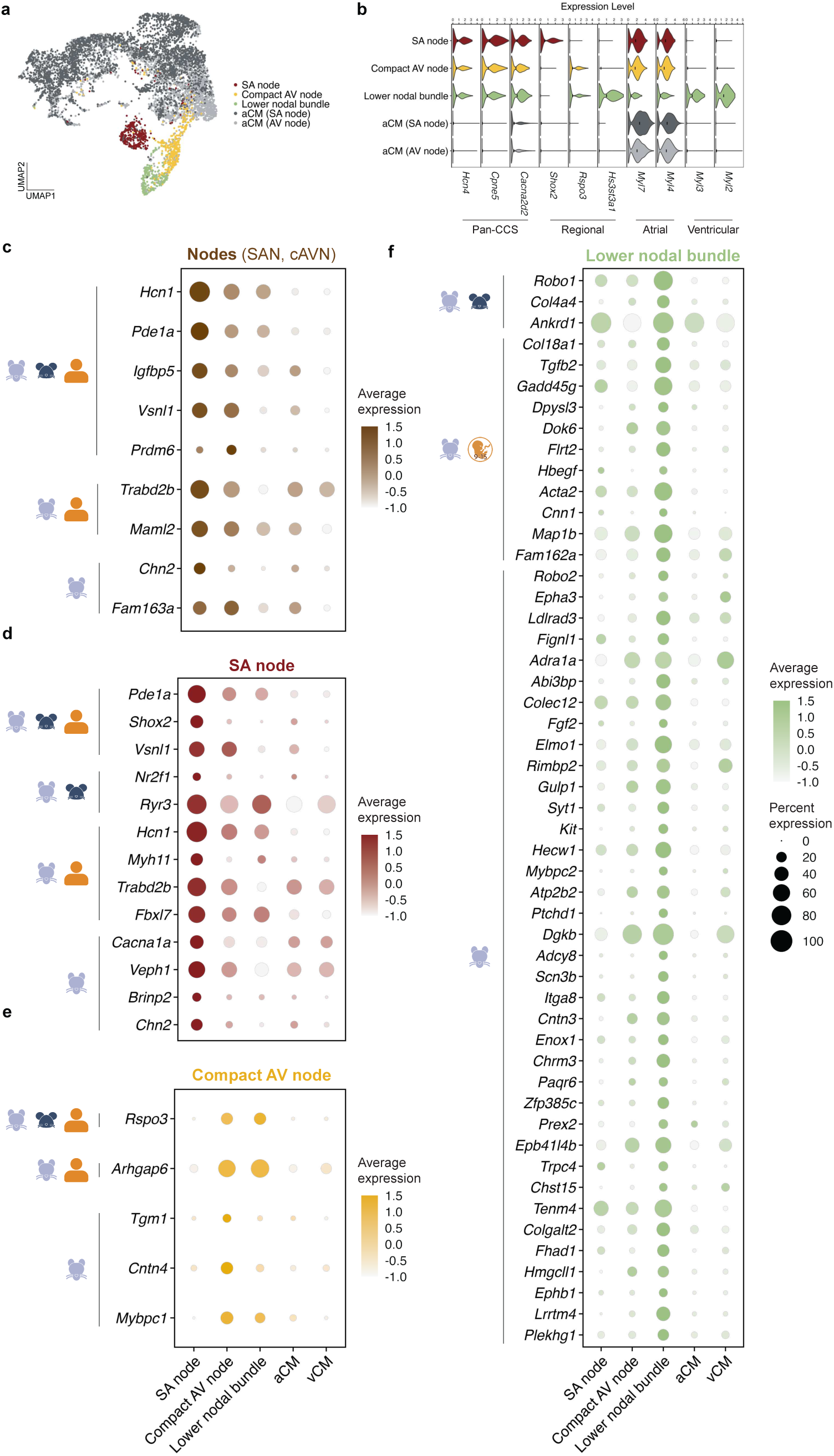
Conserved and species-specific molecular profiles in the adult rat CCS. **a**, UMAP embedding of cardiomyocytes isolated from adult rat micro-dissected SA node and AV node anatomical regions. **b**, Expression of marker genes used for cell type annotation. **c-f**, Expression of conserved and rat-specific genes enriched in the nodes (**c**), SA node (**d**), compact AV node (**e**), and lower nodal bundle (f). Rat-specific genes are defined as those expressed in <5% of the corresponding region in both adult human and postnatal mouse. Icons indicate the species in which each gene is enriched.

#### Sinoatrial node

Among CCS components, the SAN was the only region that showed conserved markers across both species and developmental stages. These included *SHOX2*, *PDE1A*, *GNAO1*, *PRDM6*, and *VSNL1* (**Fig. 3a**). Notably, *VSNL1* has recently been reported as an SAN marker in mice, zebrafish, rabbit, and cynomolgus monkey^17, 23, 48^. Spatial visualization confirmed *PRDM6* expression in the SAN region of both fetal and adult human hearts (**Fig. 3b**).

In human fetal and adult SAN, as well as in the mouse embryonic SAN, but not in the postnatal mouse SAN, *TDRP* and *CDH11*, *HCN1,* and *TENM4* were identified. Notably, key developmental TFs, *TBX18* and *ISL1*, which were undetectable in scRNA-seq or spatial transcriptomics in the postnatal mouse SAN after birth^14^, were contrarily detected in the adult human SAN, highlighting stage– and species-specific differences.

The developing SAN shared additional markers across species: *BMP4*, known to be expressed in pacemaker cells^18^, *PDGFRA*, *GATA6*, and *FZD7*, which are all primarily associated with developmental signaling. Furthermore, several axon guidance genes, such as *TENM3*, *EFNB2*, *ZDHHC2*, *CNTFR,* and *UCHL1*, were strongly conserved, suggesting an implication of neuronal genes in SAN morphogenesis. On the other hand, only one marker, *PRSS23*, which encodes serine protease 23 and is involved in cardiac valve development^48^, was conserved across species.

Additionally, we identified genes conserved as SAN markers within the same species (**Fig. 3a**). The human fetal and adult SAN expressed ion channel genes, such as *KCNJ3* (encoding acetylcholine-activated potassium channel subunit Kir3.1)^39^, *CACNA1D* (voltage-gated calcium channel subunit Ca_V_1.3)^39^, and *TRPM3* (nonselective transient receptor potential cation channel); receptors, including *PTPRD* (protein tyrosine phosphatase receptor type D), *GNAQ* (G protein subunit alpha Q), and *NPR1* (neurophillin1, a co-receptor for VEGF); cell adhesion genes, such as *CNTN4*, *FAT3* and *BOC*; as well as enzymes like *PLCB1*, *CPE*, and *ADAMTS19*. Conversely, the mouse SAN at both embryonic and postnatal stages was specifically marked by *Gria1* (glutamate ionotropic receptor), *Grip1* (glutamate receptor interacting protein 1), and *Gcgr* (glucagon receptor that modulates heart rate in mice^49^). Also, several unexplored genes, including *RGS12*, *DCTN6*, *ALOX15B*, *ADAM33*, *REC114*, and *NRSN2*, were identified as mouse SAN markers.

These stage– and species-specific patterns underscore the dynamic SAN transcriptional changes during development and maturation. To further resolve transcriptional changes in the developing SAN, we integrated a recently published scRNA-seq data of the human fetal SAN at 19 pcw^18^ with the 9-15 pcw dataset (**Supplementary** Fig. 4a). Based on strong concordance between matching cell types in the two datasets (**Supplementary** Fig. 4b), we performed a pairwise comparison of genes that were conserved in human and mouse SAN populations (**Supplementary** Fig. 4c). Notably, genes related to electrophysiological function (e.g., *HCN1*, *CACNA2D2*) and axon guidance (e.g., *TENM4*, *ZDHHC2*) increased from 9-15 pcw to 19 pcw, whereas genes involved in SAN development (e.g., *SHOX2*, *BMP4* and *FZD7*) decreased. To further contextualize these temporal patterns, we examined the expression of these genes in adult human SAN (**Supplementary** Fig. 4d) and human induced pluripotent stem cell-derived SAN (iPSC-SAN)^32^ (**Supplementary** Fig.

**4e**). In general, genes rising during fetal development maintained high expression in the adult SAN, while those with reduced expression were generally low in the adult SAN, further supporting transcriptional changes during maturation. Interestingly, most SAN genes continued to rise during iPSC-SAN differentiation, indicating that these cells are developmentally immature, possibly corresponding to a stage earlier than 15 pcw.

#### Compact atrioventricular node

In the cAVN, atrioventricular canal markers, *BMP2*^37^ and *RSPO3*^50^, were conserved in humans and mice, with *BMP2* conserved at the developing stage and *RSPO3* at the adult/postnatal stage (**Fig. 3c**). AVN-specific expression of *RSPO3* in human fetus was supported by spatial visualization, showing its expression in the AV ring as well as tricuspid valve interstitial cells (**Fig. 3d**).

Developing cAVN markers shared by both species included known CCS genes, *ID2*^46^, *CPNE5*^22^, *IGFBP5*^22^, and *CACNA2D2*^1^, as well as *KDR* (VEGF receptor associated with congenital heart disease)^51^, and *SMIM1*, neither of which had been previously linked to cAVN function. No postnatal/adult-specific cAVN markers were identified.

cAVN markers conserved in human across developmental stages included *ST6GALNAC3* (a sialyltransferase enzyme), and *TNNI1* (slow skeletal muscle isoform of troponin) (**Fig. 3c**).

Furthermore, several cAVN markers were conserved between embryonic and postnatal mice, including *Sfrp5*, *Slc22a1* (Organic cation transporter 1), *Nes*, *Nfasc*, *Slitrk5*, and *Rbpms* were identified.

#### His bundle

At the adult/postnatal stage, *FBN2* (fibrillin 2) was a conserved HIS marker between species (**Fig. 3e**). Spatial visualization verified *FBN2* expression in human fetal and adult HIS, with lower expression in AVN/atrioventricular ring and septal cardiomyocytes (**Fig. 3f**). In the developing HIS, we observed cross-species conservation of *ATP1B1* and *HS3ST3A1*, which we recently identified as LNB/Proximal VCS and cAVN/LNB markers, respectively, in the postnatal mouse CCS^14^.

Stage-shared human HIS markers included *CPMX2*, *COL5A2*, *GABRE*, *EPHB3*, and *SLC27A6*. Given recent findings showing that a GABAergic system modulates electrical conduction in mouse AVN^52^, *GABRE* (GABA receptor subunit epsilon) may also influence atrioventricular conduction in human HIS (**Fig. 3e**). In addition, long non-coding RNAs, *CRNDE* and *LINC01405* (*VHRT*) marked human HIS, while their roles in the CCS remain unknown. Conversely, in mice, both embryonic and postnatal HIS exhibited marker genes, including *Lyz2* (the mouse ortholog of *LYZ*, conventionally used as a myeloid marker^53^), *Kcne1*, *Gpr22*, as well as others recently identified and validated^14^, such as *S100a6*, *Igfbp7*, and *Ppp1r17*. Notably, these markers are not evident in human HIS.

#### Purkinje fibers

In the developing PF, several trabecular genes were conserved, including *ANGPT1*^54^, *SEMA3A*^29^, *IRX3*^28, 55^, *NPPA*^56^ and *GJA5*^3^, suggesting that PF formation is guided by evolutionarily conserved gene programs originating in trabecular myocardium (**Fig. 3g**). Additional conserved markers of the developing PF included *KCNK3* (TASK-1 potassium channel; also conserved in chicken)^57^; *SCN5A* (Na_V_1.5 sodium channel, causally associated with Brugada syndrome)^58^; and *TGM2* (transglutaminase 2). Spatial visualization showed *KCNK3* expression in the PF as well as the atria in the fetal human heart, while it was absent in the human adult PF (**Fig. 3h**).

While no genes were conserved in adult PF between species, we found human-specific PF markers, which included *NCALD* (encoding a neuronal calcium sensor protein, neurocalcin), *ABTB2,* and *LMOD2* (**Fig. 3g**). In addition, in mice, both developing and adult mouse PF were marked by *Bmp10* (ligand associated with mouse trabecular development)^50, 59^, and *Scn10a* (Na_V_1.8). The human ortholog of *Scn10a* is highly related to Brugada syndrome and other conduction abnormalities^60^, whereas *SCN10A* expression was scarce in the human CCS (**Fig. 3g**).

Our analysis also revealed unique species– and stage-specific CCS component markers (**Supplementary Table 3**) that are specific to one species or stage and absent from the other. For example, *BDNF* (brain-derived neurotrophic factor) is a specific marker for developing human SAN, while *TNNT3* (fast skeletal muscle troponin isoform) marks the developing cAVN, with neither gene expressed in the corresponding mouse components. In addition, while *Cntn2* is a well-known pan-CCS marker in mice^9, 14, 61–63^ but not humans, *CNTN4* is SAN-specific, and *CNTN5* is adult HIS-specific in humans, with neither present in the equivalent mouse components.

### Conserved and species-specific CCS transcriptional profiles in rats

Next, we extended our comparative analysis to rats, another commonly used model organism in cardiac electrophysiology, by using a recently published single-cell transcriptomic atlas that includes adult rat SAN and AVN cardiomyocytes (**Fig. 4a**)^21^. Particularly, corroborating the recent profiling of the mouse CCS^14^, the rat AVN was further subclustered into the cAVN and LNB. The LNB was defined by hybrid expression of ventricular and atrial chamber markers, *Myl2* and *Myl7*, respectively, and it expressed a newly identified marker gene, *Hs3st3a1*^14^ (**Fig. 4b**). We then derived regionally enriched gene sets, both against contractile atrial cardiomyocytes and other CCS populations, for (i) both nodes combined (SAN + cAVN), (ii) SAN only, (iii) cAVN only, and (iv) LNB only (**Supplementary Table 2**).

Comparison with adult human and postnatal mouse data revealed a core set of CCS zone and component-specific markers conserved across all three species: *Hcn1*, *Pde1a*, *Igfbp5*, *Vsnl1,* and *Prdm6* in both nodes; *Pde1a*, *Shox2* and *Vsnl1* in the SAN; and *Rspo3* in the cAVN (**Fig. 4c-e**). Additionally, we identified markers conserved in only two species, including *Nr2f1* and *Ryr3*^23, 64^ in rat and mouse SAN; *Myh11*, *Trabd2b*, and *Fbxl7* in rat and human SAN; and *Arhgap6* in rat and human cAVN.

Since human adult CCS data does not include LNB, we incorporated the human fetal dataset (9 – 15 pcw) that contains the LNB population and performed a cross-species comparison (**Fig. 4f**). LNB markers shared between rat and mouse included *Robo1*, *Col4a4,* and *Ankrd1*. Rat and human LNB also shared *Flrt2*, *Acta2*, and *Cnn1*. Notably, *Tgfb2* and *Hbegf*, required for atrioventricular cushion remodelling, septation, and cardiac valve formation in mice^65, 66^, were conserved between rat and human LNB as well.

Additionally, rats displayed species-specific CCS markers, which were expressed in less than 5% of cells in the corresponding regions of the other datasets. In the SAN, *Cacna1a* (neuronal calcium channel Ca_V_2.1) was uniquely enriched (**Fig. 4d**). The rat cAVN expressed *Cntn4* as well as *Mybpc1* (slow skeletal paralog of myosin-binding protein C) (**Fig. 4e**). In the LNB, rat-specific genes included *Robo2* (a family member of the proximal VCS marker *Robo1*); *Mybpc2* (fast skeletal paralog of myosin-binding protein C); *Cntn3* (contactin 3); *Scn3b* (sodium channel β-subunit); and *Tenm4* (axon guidance protein) (**Fig. 4f**). Together, these results demonstrate CCS-region specific transcriptomic heterogeneities with clear species-specific variations.

### Conserved and species-specific CCS transcriptional profiles in zebrafish and medaka

As an emerging model in cardiac electrophysiology, zebrafish is known to express a number of conserved CCS genes^67^. However, direct comparisons between the developing and mature zebrafish CCS, as well as between zebrafish and other mammalian CCS at a single-cell level, have not been conducted. To address this, we utilized scRNA-seq datasets from both embryonic^17^ and adult^16^ zebrafish, containing SAN and atrioventricular canal (AVC) populations, to identify CCS markers and compared them with SAN and cAVN/AVC gene expression in humans and mice (**Fig. 5a-b** and **Supplementary Table 2**).

**Figure 5.**
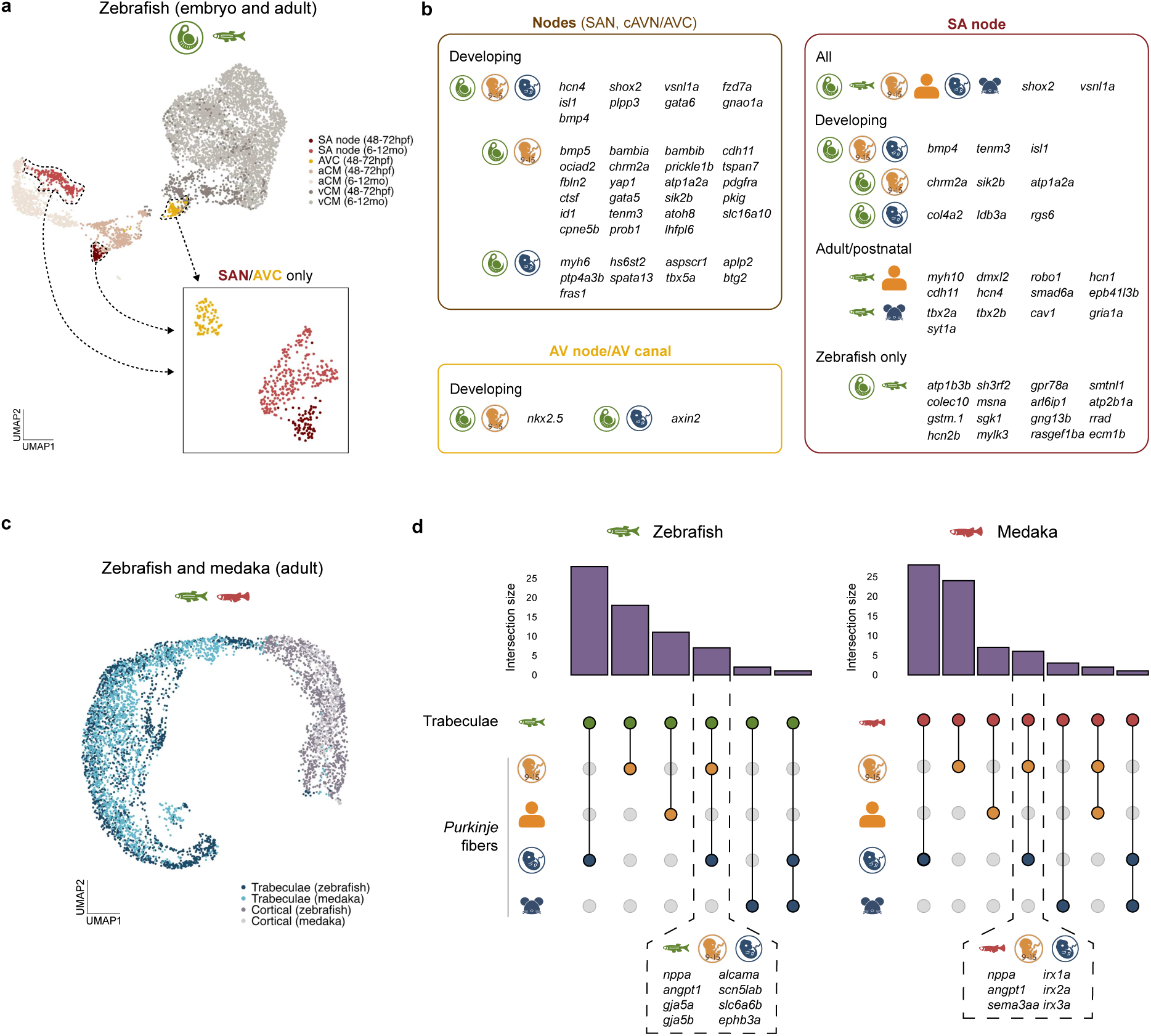
Conserved and species-specific molecular profiles in the zebrafish and medaka CCS. **a**, Integrated UMAP embedding of embryonic and adult zebrafish cardiomyocytes. Inset shows integrated UMAP embedding excluding working cardiomyocytes. **b**, Genes enriched in the SA node and/or AVC, grouped by conservation across species and developmental stages. **c**, Integrated UMAP embedding of adult zebrafish and medaka ventricular cardiomyocytes. **d**, Shared markers between zebrafish and medaka trabeculae and mammalian *Purkinje* fibers. Bars represent the number of genes in each intersection (based on human orthologs). Insets highlight genes conserved across all three species. Zebrafish and medaka trabeculae markers were defined relative to cortical myocardium, and *Purkinje* fiber markers were defined relative to both other CCS cell types and to surrounding ventricular cardiomyocytes.

Among nodal markers, *vsnl1a* was the only gene conserved across all species and stages, extending previous findings of broad conservation of *Vsnl1* in mammals^23^ to non-mammalian vertebrates as well. Several TFs were also broadly conserved, including *shox2*, *gata6*, and *isl1* across developing zebrafish, human, and mouse nodes; *id1*, *atoh8* and *gata5* between zebrafish and human; and *tbx5a* between zebrafish and mouse. We also observed conserved expression of signaling genes, such as the Wnt receptor *fzd7a*, conserved in zebrafish, human, and mouse, and *bmp4*, conserved in zebrafish and human.

In the adult/postnatal nodes, we observed conservation across all three species of *hcn1*, encoding a pacemaker channel. We also observed conservation of several TFs, including *smad6a*, conserved in all three species, *znf536*, conserved in zebrafish and human, and *tbx2a*, *tbx2b* and *gata5*, conserved in zebrafish and mouse. Notably, *robo1*, a well-known proximal VCS marker in mice^26, 27^, was conserved as a nodal marker in adult zebrafish and humans.

Many genes were conserved only between embryonic and adult zebrafish, suggesting species-specific nodal programs. These included *cx36.7*, a slow-conducting gap junction orthologous to mouse Cx30.2; *colec10* and *atp1b3b*, which have been shown to affect heart rate in embryonic zebrafish^17^; *vcana* and *bcam*, orthologs of conserved VCS-enriched genes in humans and mice (**Fig. 2b**); and the Wnt ligand *wnt2bb*. Direct comparison of embryonic and adult zebrafish SAN revealed distinct stage-dependent expression patterns, with both stages showing enrichment of different functional gene categories, including ion channels, sarcomeric components, ligands, and TFs (**Supplementary** Fig. 5). Notably, many TF genes (e.g., *gata6*, *isl1*, *tbx2a*, *tbx2b*) were enriched in the embryonic SAN.

Unlike mammals, teleost fish do not possess a morphologically distinct VCS^68^. Instead, trabecular cardiomyocytes, which comprise most ventricular cardiomyocytes^69^, have been proposed to support rapid impulse propagation and synchronized ventricular activation, making them functionally analogous to mammalian embryonic trabeculae and PF^70^. To compare the molecular profiles of these cell populations, we used scRNA-seq data containing cortical and trabecular cardiomyocytes from adult zebrafish and medaka^15^. We identified genes enriched in trabecular versus cortical cardiomyocytes in each fish species (**Fig. 5c**) and assessed overlap with genes enriched in mammalian PF, compared to surrounding non-CCS cardiomyocytes as well as other CCS components (**Fig. 5d**). Although the fish data are from adult hearts, we included both embryonic and adult mammalian PF populations in our comparison, as PF originate from embryonic trabeculae^71^.

Across all four species – embryonic mouse, fetal human (9 – 15 pcw), adult zebrafish, and adult medaka – we observed conserved expression of the signaling molecules *nppa* and *angpt1*. In zebrafish, additional genes were conserved with embryonic mouse and fetal human PF, including the ephrin receptor *ephb3a*, both zebrafish orthologs (*gja5a* and *gja5b*) of the mammalian fast-conducting gap junction *GJA5*, and *scn5lab*, an ortholog of the cardiac sodium channel *SCN5A*. These were likely not detected in medaka due to incomplete genome annotation. In medaka, we observed conserved expression of the semaphorin ligand *sema3a*, as well as the *Iroquois* homeobox TFs, *irx1a*, *irx2a*, and *irx3a*, each shared with embryonic mouse and human PF.

### Conserved transcriptional regulation of CCS heterogeneity

Although several individual transcription factors have been extensively studied, primarily in the mouse CCS, the transcriptional regulatory networks of the entire human CCS components, and their conservation relative to the mouse, remain to be further defined. To address this gap, we applied SCENIC (single-cell regulatory network inference and clustering)^72, 73^ to single-cell transcriptomes of fetal and adult human CCS as well as embryonic mouse CCS, systematically mapping TF activities and their regional specificity (**Supplementary Table 4**, as previously performed^14^. Using Pearson correlation of TF activity scores, we identified regulators with similar activity patterns between human and mouse CCS components at each developmental stage (**Fig. 6a-b**).

**Figure 6.**
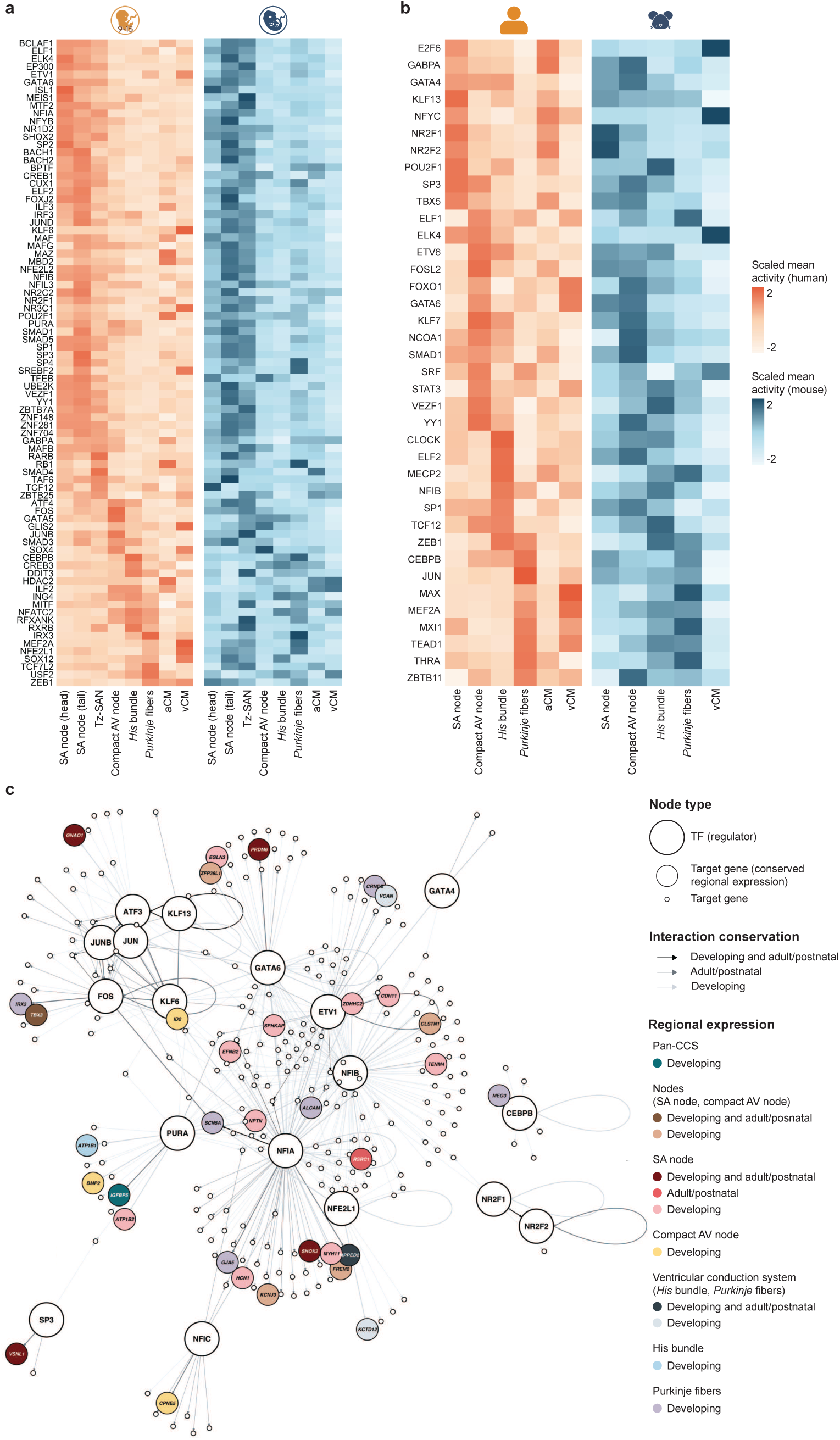
Conserved transcriptional regulation of CCS heterogeneity. **a-b**, Activities of conserved regulons across CCS components in fetal (**a**) and adult/postnatal (**b**) human and mouse hearts, quantified by area under the recovery curve (AUC) of target genes inferred using the SCENIC algorithm. Only activating regulons are shown. **c**, Gene regulatory networks depicting transcription factor (TF)-target gene interactions conserved between human and mouse CCS.

In the fetal CCS, several TFs exhibited regionally clustered activities, revealing both component-specific and broadly acting regulators in humans and mice (**Fig. 6a**). These include well-established CCS TFs, such as GATA6, ISL1 and SHOX2 in the SAN^4, 5, 34^ and IRX3 in HIS and PF^10, 11^. We also identified TFs not previously well-linked to the CCS, such as ELF1, NFIA, and NR2F1 in the SAN; GATA5 and SOX4, known to regulate gap junctions with TBX3^74^, in cAVN; CEBPB and CREB3 in HIS; and ING4 in HIS and PF. Similarly, the adult/postnatal CCS displayed region-specific TF activities, such as KLF13, NR2F1, NR2F2 in SAN; SP3 and TBX5 in SAN and cAVN; KLF7, NCOA1, and YY1 in cAVN; NFIB in HIS; TCF12 in cAVN and HIS; ZEB1 in HIS and PF; and TEAD1 in PF (**Fig. 6b**).

Then, to examine conserved transcriptional regulation of CCS genes between humans and mice, SCENIC predicted TF-target interactions were obtained from each dataset and compared between the fetal human and embryonic mouse CCS and between the adult human and postnatal mouse CCS. Based on conserved TF-target gene interactions, we reconstructed conserved gene regulatory networks for the CCS (**Fig. 6c**). Consistent with previous studies on ETV1 in the VCS^9^ and atrial conduction^75^, our results indicated that ETV1 regulates both SAN genes (e.g., *SPNKAP*, *CDH11*) and PF genes (e.g., *ALCAM*). It is also notable that FOS, JUN, JUNB, ATF3 and KLF6, which are all bZIP proteins in the AP-1 network and physically dimerize with specific partners, cluster together in regulating CCS markers, such as *IRX3*, *TBX3*, *ID2* and *GNAO1*.

Together, these analyses predict conserved gene regulatory mechanisms across species, which underlie transcriptional heterogeneities within the CCS.

### Disease associations of conserved regional markers

Lastly, we assessed the functional and clinical relevance of CCS markers identified in this study, encompassing region-enriched genes conserved across species, as well as genes conserved across developmental stages in humans. Specifically, we conducted the association analyses of common and rare genetic variants in these genes with cardiac conduction-related traits and diseases, categorized into groups, such as atrial conduction (atrial arrhythmias/P wave), AV nodal function (PR interval/segment), and other conduction phenotypes.

Analysis of common genetic variants from genome-wide association studies (GWAS) for 31 cardiac conduction-related traits revealed significant associations for 57 genes (**Fig. 7a** and **Supplementary Table 5**). In addition to well-characterized CCS genes (e.g., *HCN4*, *SCN5A*), we uncovered genes with no prior characterization in the CCS. For example, SAN-enriched *CDH11* is associated with heart rate; LNB-enriched *HBEGF* is associated with PR interval; and HIS-enriched *SLC27A6* is associated with PR interval and ventricular arrhythmia.

**Figure 7.**
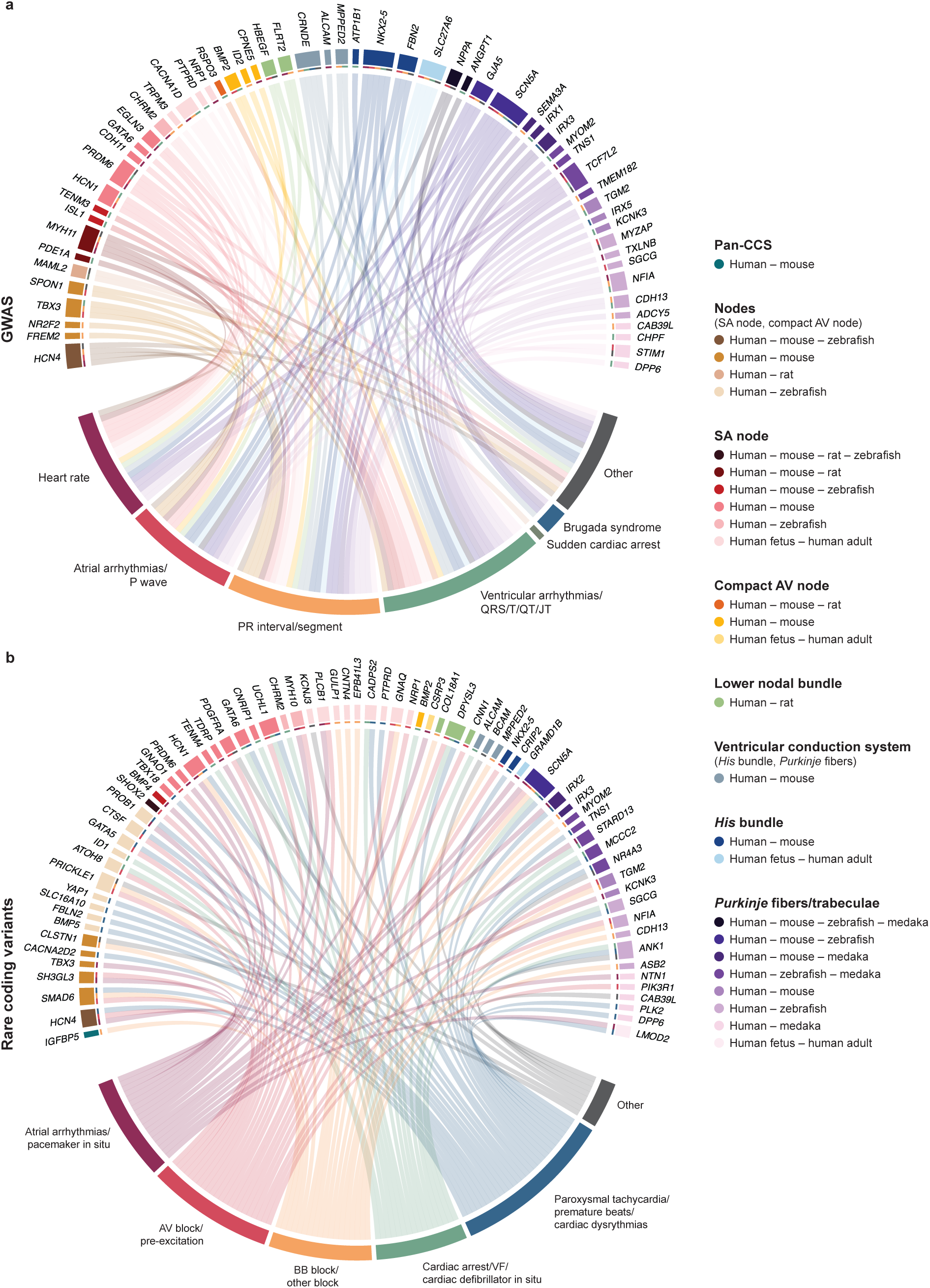
Cardiac conduction-related functional and disease associations of conserved CCS markers. **a-b**, Circos plots showing associations of species– and stage-shared CCS genes, grouped by regional enrichment, with electrophysiological traits and conduction diseases. Associations were identified via the GWAS catalog (**a**) and phenome-wide analysis of rare coding variants (**b**). When applicable, species conservation is prioritized over stage conservation for labeling. Genes enriched in multiple components as well as the encompassing broader zone are labeled by the broader zone designation. AV block, atrioventricular block; BB, block bundle branch block; VF, ventricular fibrillation.

Moreover, we examined whether rare coding variants, computationally predicted to be deleterious, in our gene set were linked to cardiac conduction diseases. Using a recently published dataset^76^, 69 genes with significant associations with conduction phenotypes in humans were identified (**Fig. 7b** and **Supplementary Table 5)**. As expected, node-enriched ion channel genes, such as *HCN4*, *HCN1*, and *CACNA1D*, were linked to heart rate and/or atrial arrhythmias, consistent with their roles in pacemaker activity. Notably, rare coding variants in node-enriched *ATOH8* and *GATA5*, as well as SAN-enriched *GNAO1* and *UCHL1*, genes which do not have a known role in the CCS, were associated with AV block. Similarly, coding variants VCS-enriched *ALCAM* and *MPPED2*, as well as PF-enriched *TGM2* and *NFIA*, were linked to AV block and/or bundle branch block.

It should be noted that while many conserved CCS genes demonstrated associations consistent with the function of their corresponding regions, other associations did not align directly with expected regional roles. This suggests that CCS gene-trait relationships may extend beyond canonical regional functions or may reflect the tissue complexity underlying arrhythmias.

Collectively, these findings highlight evolutionarily conserved, region-specific CCS genes likely with functional and pathological importance in humans, expanding the pool of candidate genes for conduction disorders.

## DISCUSSION

In the present study, we systematically compared single-cell and single-nucleus transcriptomic data from CCS components (or equivalents) of humans, mice, rats, zebrafish, and medaka across developmental and mature stages. To facilitate data sharing with the research community, we have developed an easily accessible web application (https://ccsatlas.com), which enables detailed, stage– and species-specific exploration of CCS gene expression.

Our comparative analysis uncovered evolutionarily conserved, regional-specific gene programs that underpin core aspects of CCS development and function. The conservation of well-known TFs, such as *SHOX2*, *TBX3,* and *GATA6* in the nodes, and *IRX3* in the VCS, reinforces the importance of shared gene regulatory networks in establishing CCS heterogeneity, as we have recently demonstrated in postnatal mice^14^. This finding is further corroborated by the conservation of developmental signaling genes, including TGF-β signaling ligands, *BMP2* and *BMP4*, ephrin signaling genes, *EFNB2* (in the nodes) and *EPHB3* (in VCS) and the Wnt receptor *FZD7*, although the roles of several conserved genes remain to be elucidated. A notable subset of the conserved CCS markers included neuronal genes. Examples include the axon guidance genes, *TENM3* and *TENM4* in the nodes and neural adhesion genes, *NPTN* and *ALCAM* in the nodes and VCS, respectively.

Importantly, genetic variants in many of these conserved CCS markers, including neuronal genes, are significantly associated with arrhythmias and other conduction disorders, supporting their functional importance and translational capacity for human conduction disorders. Because the roles of many conserved genes in the CCS development, function and disease are largely unknown, future studies are warranted.

Despite broad conservation, certain genes exhibited species-specific expression, which may reflect physiological differences. Notably, transcriptional similarity across species was higher in SAN and nodal regions, whereas greater divergence was observed in the VCS and PF. This likely reflects the universally conserved role of the SAN as the impulse generator, in contrast to PF, which varies in cell size, fiber thickness, action potential profiles, conduction velocity, as well as structural organization, particularly between rodents and primates^77–79^.

Findings from our comparative analysis have several implications. First, many CCS-enriched genes are not currently included in gene panels for inherited arrhythmias. This highlights a potential diagnostic gap for conduction disorders, as a significant number of arrhythmias lack an identifiable genetic cause. Thus, inclusion of conserved CCS genes which are further functionally validated can improve genetic risk prediction and management for individuals living with conduction disorders.

Second, as the progress and development of single-cell omics continue, functional validations using preclinical models will become essential. Knowing which genes are conserved or divergent will help researchers choose the most appropriate model organism. For example, TBX18 has been used to drive direct SAN differentiation *in vivo* by promoting pacemaker activity^80, 81^. Our analysis revealed that while it is undetectable in the mouse SAN after birth, it is persistently expressed at low levels in the adult rat SAN and adult human SAN, suggesting it may support adult SAN function in humans and rats. Consequently, TBX18-based direct reprogramming strategies might yield species-specific outcomes.

Lastly, our findings support stem cell-based regenerative medicine. Generating biological pacemaker cells using iPSC-based differentiation has been proposed as a promising therapeutic approach^18, 81^, but the SAN’s cellular and genetic heterogeneity implies that more diverse reprogramming methods may be required. Likewise, iPSC differentiations toward cAVN, HIS, and PF remain to be further investigated and developed. Therefore, defining gene regulatory networks for each CCS component will guide the refinement of differentiation protocols.

Nevertheless, our study has several limitations. Firstly, the low number of cells for some CCS components, such as adult human PF cells and mouse postnatal SAN cells, as well as the low sequencing depth of sc/snRNA-seq, likely reduced the statistical power of our analysis. Due to technical variations among datasets, such as species, modality (scRNA-seq vs. snRNA-seq), and other batch effects, direct integration was not feasible. As a result, our identification of species-or stage-specific differences was also restricted to genes exclusively expressed in one species or stage, rather than differences in expression level. Our stage comparison was also limited by the imperfect matching of developmental stages. For example, due to an absence of transcriptomic data from adult mouse CCS, we considered postnatal mouse data for comparisons with adult human data. In addition, the rat, zebrafish, and medaka datasets did not include all CCS components, which precluded a full analysis of transcriptional heterogeneity across the entire CCS in these species. In addition, incomplete genome annotation and cross-species ortholog mapping, especially in zebrafish and medaka, may have obscured additional conserved genes.

In conclusion, our study provides a comprehensive, comparative transcriptomic analysis that delineates both conserved and divergent gene programs across human and mouse CCS, with supporting insights from rats, zebrafish, and medaka. By defining region-specific genes and regulatory pathways with translational relevance, our findings lay a foundation for future investigations into CCS physiology and pathology and inform efforts for the generation of CCS-like cells for disease modeling and regenerative strategies.

## METHODS

### Ethics

This study involved only secondary analysis of previously published sc/snRNA-seq datasets derived from human and animal tissues. All original studies obtained ethical approval from the appropriate institutional review boards or regulatory authorities, and details are provided in the corresponding publications. No new experiments involving human participants or animals were conducted by the authors.

### sc/snRNA-seq data processing and analysis

All sc/snRNA-seq data were processed and analyzed using Seurat^82^ (version 5.0.3). Quality control was performed as described in the original publications, with the exception of the adult human^20^ and embryonic zebrafish^17^ datasets. For adult humans, we excluded cells exceeding 30,000 detected transcripts (nCount_RNA > 30000), and for embryonic zebrafish, we excluded cells with fewer than 2,000 detected transcripts (nCount_RNA < 2000) (in addition to other quality control parameters described in the original publications). Differential expression analysis was carried out using the Wilcoxon rank sum test (*FindMarkers* or *FindAllMarkers*), with p-values adjusted for multiple comparisons using the Bonferroni correction. Parameters used in all analyses are available in the uploaded code.

CCS markers were identified using the following criteria: Genes were considered pan-CCS markers if they were differentially expressed in atrial or ventricular CCS components relative to their chamber-matched working cardiomyocytes (*P*_adj_ < 0.05, fold change > 3) and were expressed in at least 25% of cells in each CCS region. For the mouse postnatal dataset, which lacked atrial cardiomyocytes, we used P1, P2, and P4 atrial cardiomyocytes from a separate dataset^50^ (not including SAN cells) and excluded genes expressed in more than 30% of these cells. Markers for the two CCS zones, nodes (SAN and cAVN) and VCS (HIS and PF), as well as for individual components, were defined by comparing the zone of interest to the other CCS components as well as compared to non-CCS cardiomyocytes (*P*_adj_ < 0.05, fold change > 1.5, percent expression in zone of interest > 10). Where possible, non-CCS cardiomyocyte controls were matched to the anatomical location of the CCS component being analyzed. Species-or stage-specific regional markers were additional criteria expressed in less than 5% of cells in the corresponding region of the other species or stage. Species-or stage-specific regional markers (**Supplementary Table 3**) were further defined as those expressed in less than 5% of cells in the corresponding region of the other species or stage.

To integrate the two human fetal heart datasets (9 – 15 pcw^19^ and 19 pcw^18^), count matrices were merged, log-normalized and highly variable genes were selected. Expression matrices were then scaled and subjected to principal component analysis (PCA). Integration was then performed using reciprocal PCA (*IntegrateLayers*), where each dataset is projected into the PCA space of the other.

Integrated clustering was then performed by computing a shared nearest neighbour graph (*FindNeighbors)* and a uniform manifold approximation and projection (UMAP) embedding based on the reciprocal PCA.

### Orthology mapping

Orthology mapping was performed using curated tables from public databases. For human, mouse, and rat comparisons, we used the JAX orthology file (http://www.informatics.jax.org/downloads/reports/HOM_MouseHumanSequence.rpt). For zebrafish and medaka, we used ortholog mappings from ZFIN (https://zfin.org/downloads/human_orthos.txt).

Mouse orthologs for zebrafish/medaka genes were inferred indirectly through shared human orthologs, due to more complete annotations in human datasets.

### Gene regulatory network inference

Gene regulatory network inference was performed using pySCENIC^72, 73^ (version 0.12.1) as previously described. Only activating regulons were included. Regulon activity scores were calculated for each cell using AUCell and then averaged across each component. We retained only regulons in which the transcription factor was expressed in at least 10% of cells in a CCS region, was present in both species for the stage in question, and exhibited a Pearson correlation > 0 between its AUC levels in human and mouse CCS. To reconstruct conserved gene regulatory networks, we identified TF–target interactions conserved between species at either the developing or adult stage (or both), and included only CCS-specific target genes.

### Correlation analysis and hierarchical clustering

To calculate correlations between CCS cell types across datasets, count matrices were subsetted to include only CCS cell types and retained only one-to-one orthologs detected in all datasets. Each matrix was then independently normalized and scaled, and pseudobulk matrices were generated by averaging scaled expression values within each CCS cell type. Pearson correlations were then calculated pairwise between each CCS cell type (stats, version 3.6.2). The resulting correlation matrix was converted to a distance matrix (1 – correlation) and used for hierarchical clustering with average linkage (stats, version 3.6.2).

### Spatial transcriptomics

Spatial transcriptomic datasets included Visium slides from adult human hearts and MERFISH imaging from fetal heart sections. For MERFISH, we also included imputed expression values for genes not originally targeted by the probe panel (see original study for details)^19^. Genes whose imputed spatial expression was used in this study include *PDE1A*, *GNAO1*, *IRX5*, *PRDM6*, *RSPO3*, *FBN2*, and *KCNK3*. Spatial visualizations were generated using Scanpy^83^ (version 1.9.3).

### Disease association (genome-wide association studies and rare coding variants)

For common disease associations, summary statistics were downloaded from the GWAS catalog for 31 traits related to cardiac conduction (see **Supplementary Table 5**). Gene lists from each cluster were overlapped with the reported or mapped genes that had significant associations with the disease of interest in the summary statistics. This was done for genes conserved between species and between human developmental stages. For rare coding gene-disease associations, we leveraged a newly published and publicly available dataset^76^ which performed a phenome-wide association study testing the association of genome-wide rare coding variants with 601 disease phenotypes in 784,879 individuals, including over 150,000 non-Europeans. We narrowed our analysis to cardiac conduction disease-related phenotypes under the “circulatory system” category, with a nominal significance threshold of *P* < 0.05. A table containing all gene-disease associations with effect sizes and *P* values can be found in **Supplementary Table 5**. Results were visualized using ‘circlize’ in R (version 4.2.3).

## Supporting information

Supplementary Table 1: Summary of datasets

Supplementary Table 2: Species– and stage-shared markers

Supplementary Table 3: Species– and stage-specific markers

Supplementary Table 4: SCENIC-inferred regulons

Supplementary Table 5: Genetic variant associations

## ACKNOWLEDGMENTS

We thank Jarno Van der Kolk at the University of Ottawa for support in establishing a Shiny server through the Digital Research Alliance of Canada.

## CODE AVAILABILITY

All codes used to generate the results and figures in this study are available on GitHub (https://github.com/hankimlab/2025-Comparative-CCS). Other codes are available from the authors upon request.

## DATA AVAILABILITY

All sc/snRNA-seq datasets analyzed in this study were previously published and are publicly available. Dataset details are available in **Supplementary Table 1**.

## CONTRIBUTIONS

M.B., Y.O., and K-H.K. conceived and designed the project. M.B., S.D., and Y.O. conducted bioinformatic analysis and prepared figures. G.T. provided technical and academic support. Y.O. and K-H.K. directed the project. M.B., Y.O., and K-H.K. drafted, edited, and revised the manuscript with input from all authors. All authors, M.B., Y.O., S.D., G.T., and K-H.K., read and approved the final version of the manuscript.

---

**Supplementary Figure 1.**
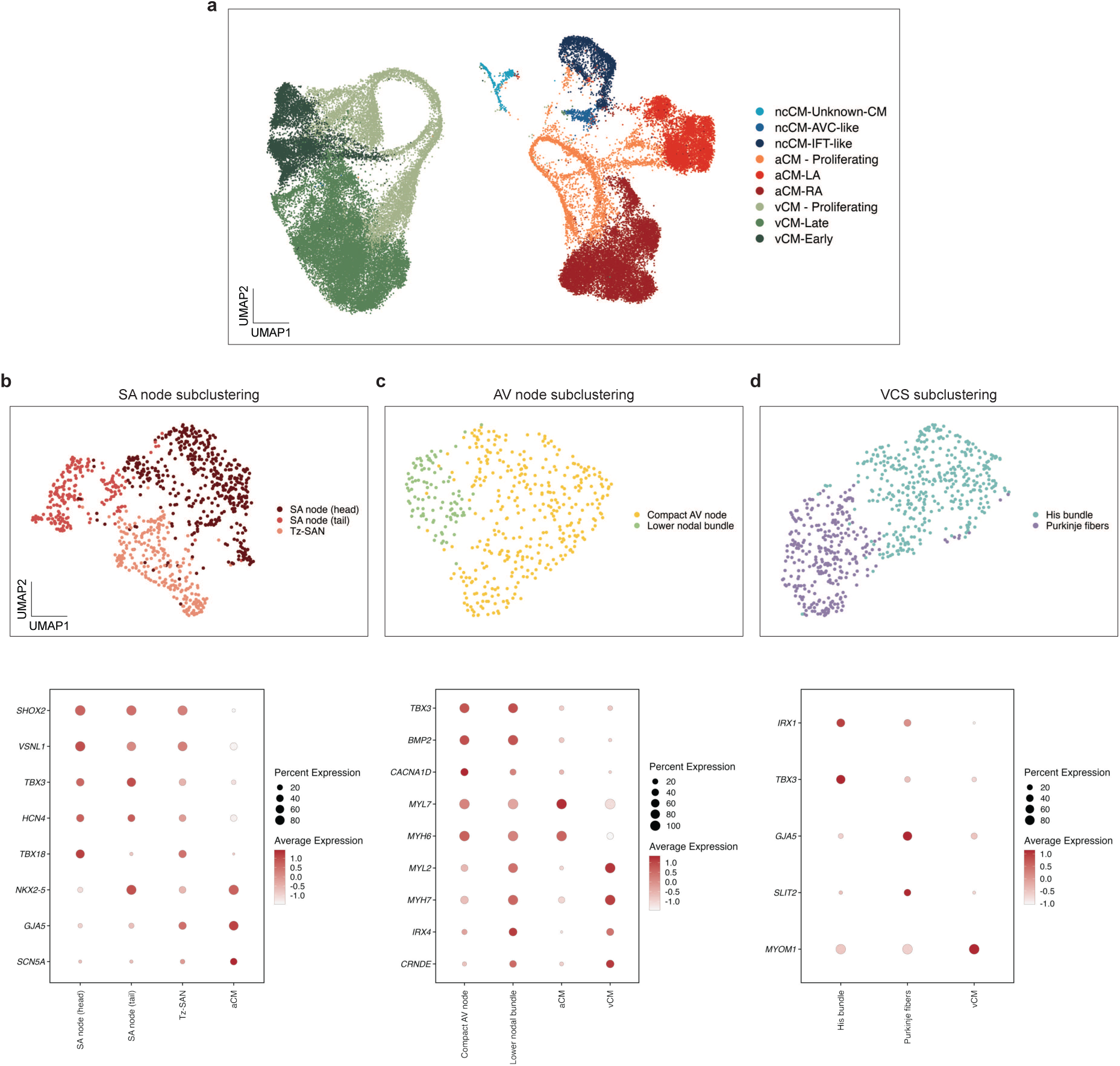
Identification of cardiac conduction system cell populations in human fetal heart single-cell RNA sequencing (scRNA-seq) data. **a**, Uniform manifold approximation and projection (UMAP) embedding of published cell annotations from human 9-15 post-conception weeks (pcw) heart scRNA-seq data (Farah *et al*, *Nature*, 2024). **b**, Subclustering of sinoatrial (SA) node cells and expression of marker genes. **c**, Subclustering of atrioventricular (AV) node cells and expression of marker genes in AV node subpopulations, atrial cardiomyocytes and ventricular cardiomyocytes. **d**, Subclustering of ventricular conduction system cells and expression of marker genes. AVC, atrioventricular canal; IFT, inflow tract; LA, left atrium; RA, right atrium; aCM, atrial cardiomyocyte; vCM, ventricular cardiomyocyte; Tz-SAN, transitional SA node; ncCM, non-chambered cardiomyocytes.

**Supplementary Figure 2.**
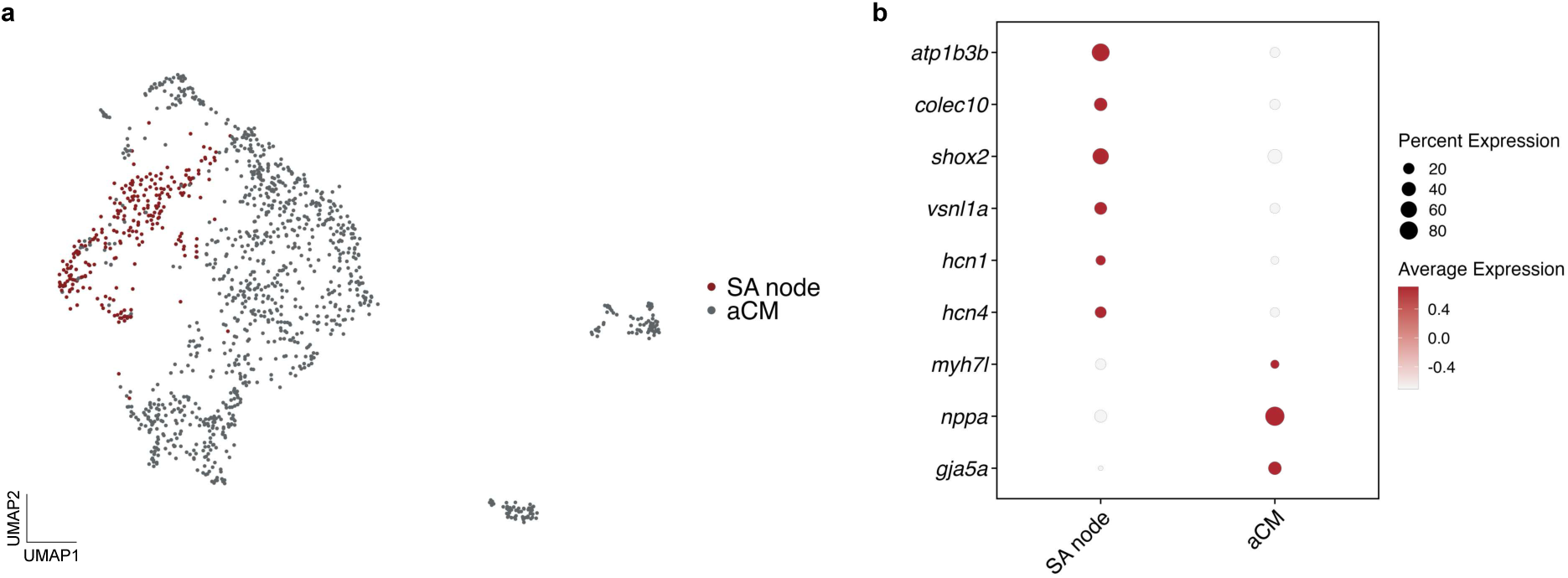
Identification of CCS cell populations in adult zebrafish heart. **a**, UMAP embedding of atrial cardiomyocytes (aCM) and SA node. **b**, Expression of canonical SAN marker genes used for cell annotation.

**Supplementary Figure 3.**
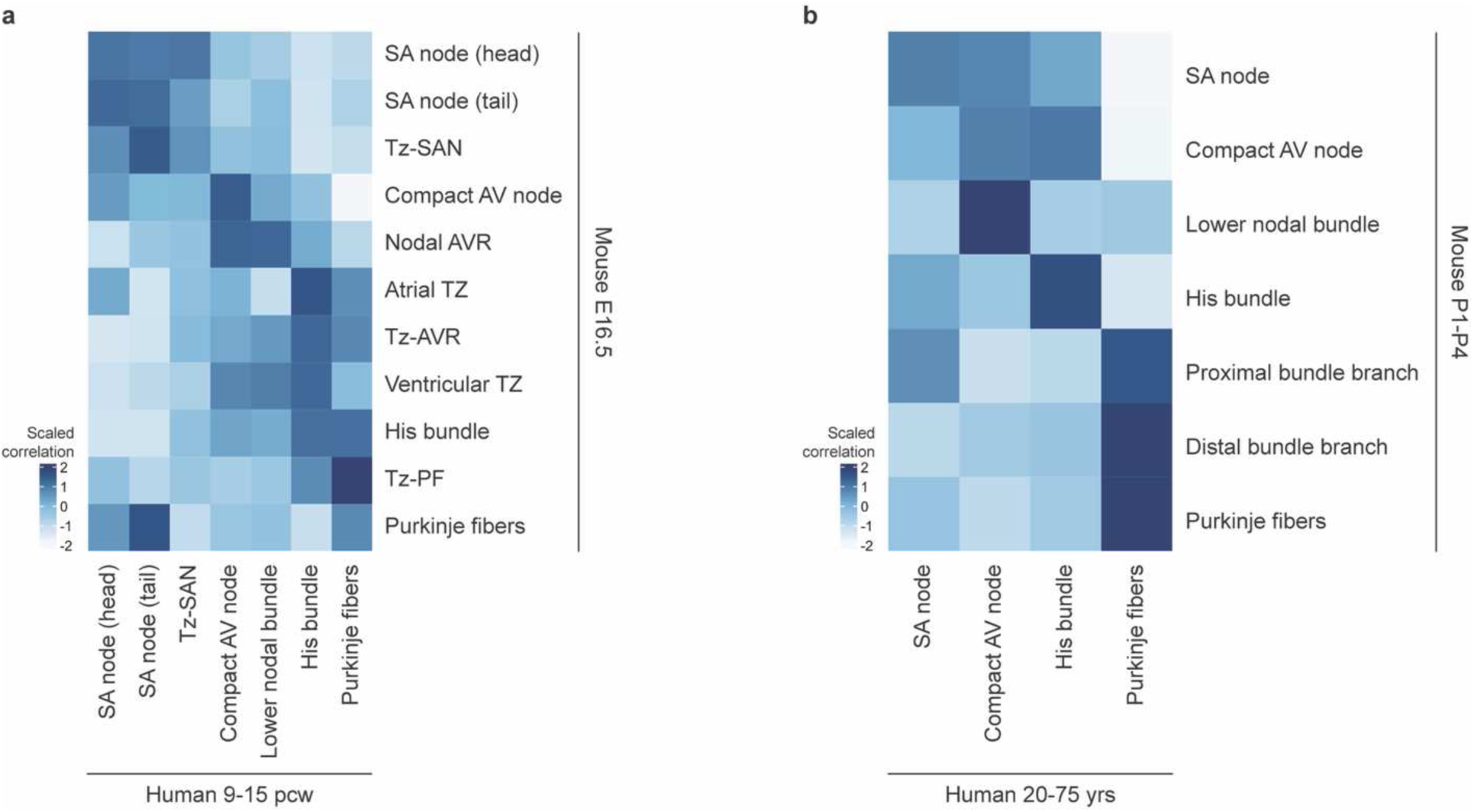
Correlations between human and mouse CCS components. **a-b**, Scaled Pearson correlation heatmap comparing gene expression profiles of CCS components between humans and mice at developmental (**a**) and adult/postnatal (**b**) stages. E16.5, embryonic day 16.5; pcw, post-conceptional week; P1-P4, postnatal days 1-4; yrs, years.

**Supplementary Figure 4.**
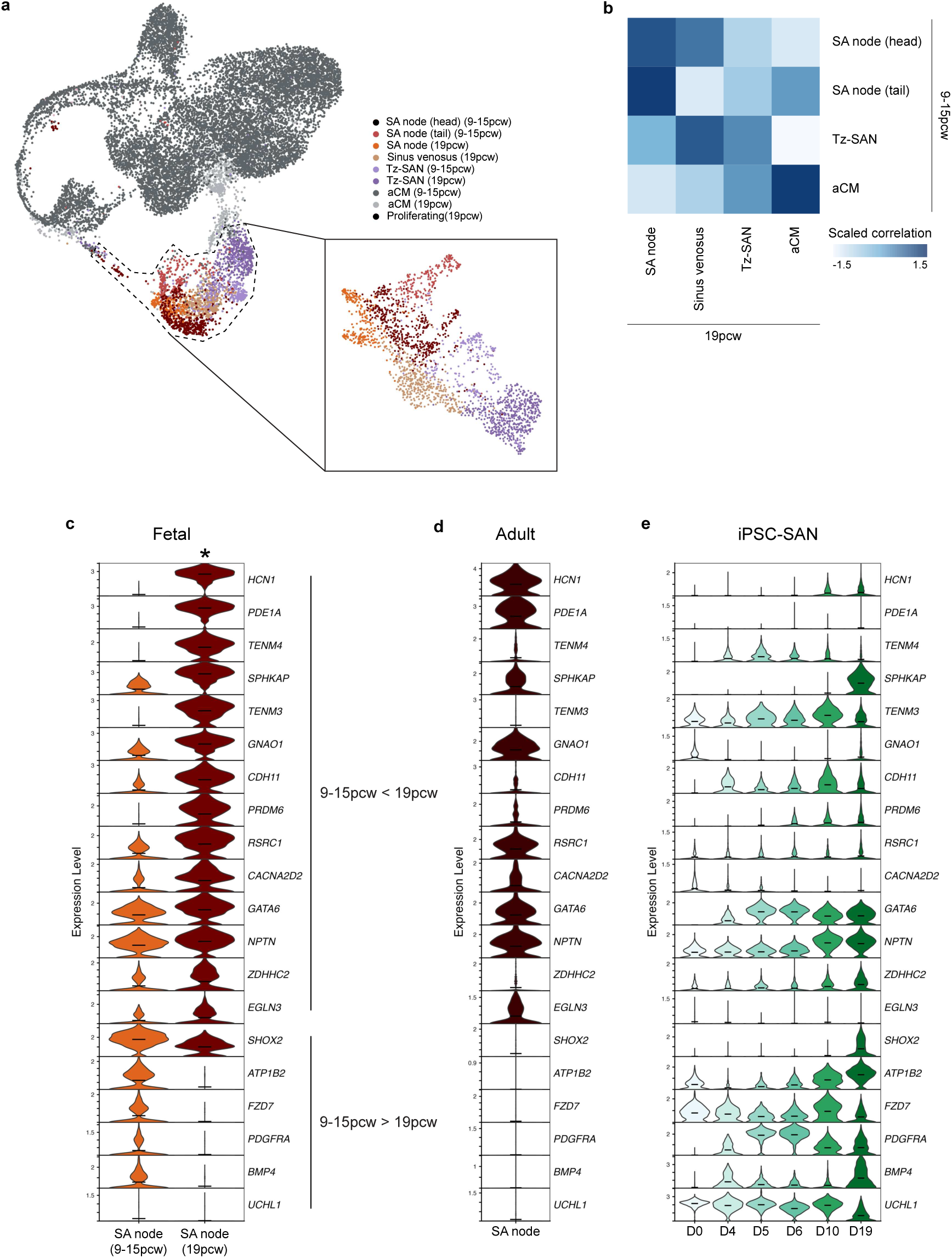
Molecular comparison of human SA node between 9-15 pcw and 19 pcw. **a**, Integrated UMAP embedding of SA node and atrial cardiomyocyte (aCM) populations from 9-15 pcw and 19 pcw fetal heart datasets. Inset shows integrated UMAP embedding excluding working aCM from both datasets. **b**, Scaled Pearson correlations between gene expression profiles of 9-15 pcw and 19 pcw cell populations. **c**, Genes conserved between human and mouse (see Fig. 2-3) that are significantly differentially expressed between the 9-15 pcw (head and tail) and 19 pcw SA node. **d-e**, Expression of the same genes in adult human SA node (**d**) and during human-induced pluripotent stem cell-derived SA node (iPSC-SAN) differentiation (**e**). *, adjusted *P* < 0.05.

**Supplementary Figure 5.**
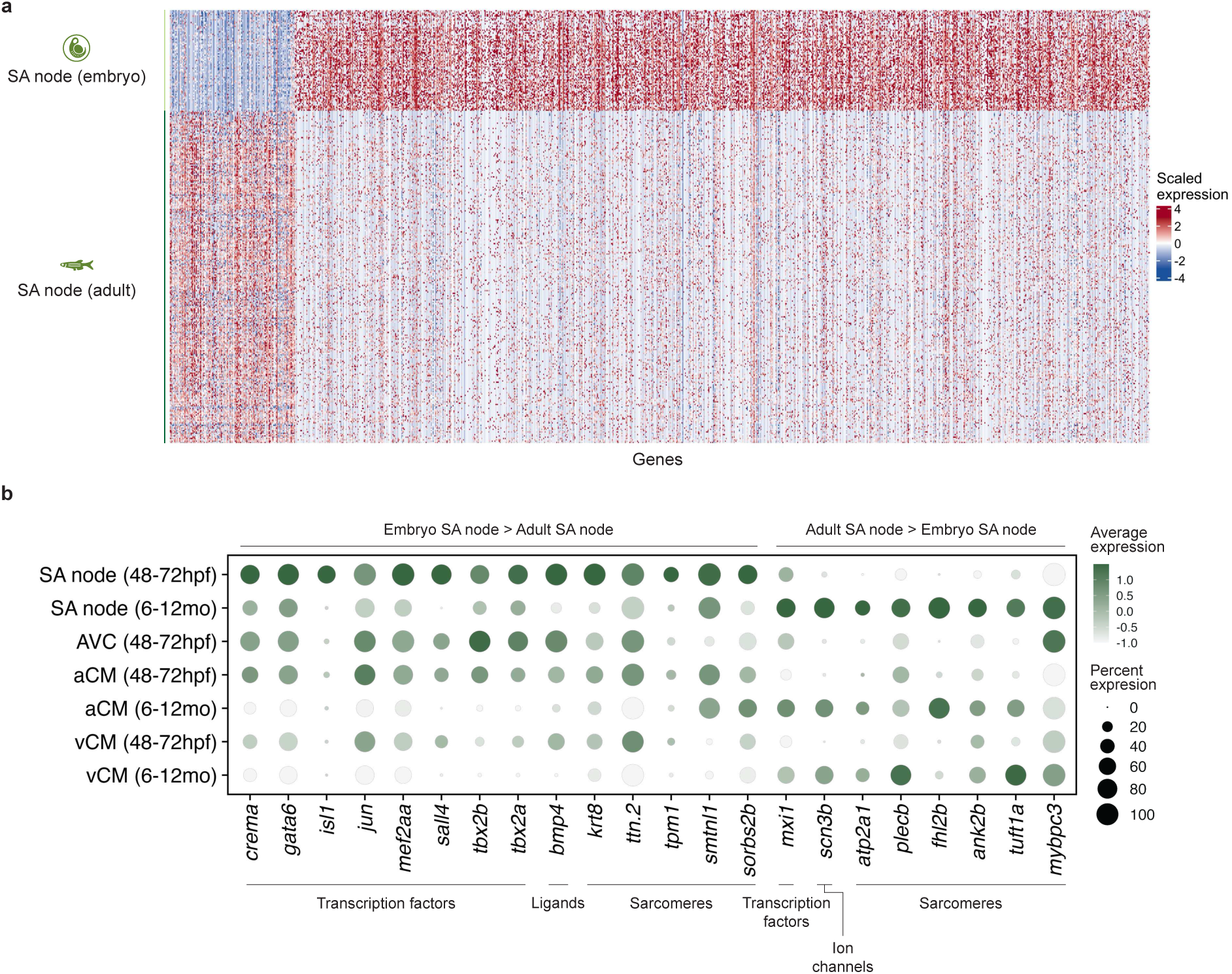
Molecular comparison of zebrafish embryonic and adult SA node. **a**, Expression of all genes differentially expressed between embryonic and adult zebrafish SA node. **b**, Expression of selected functional categories, including transcription factors, ligands, sarcomeres, and ion channels, differentially expressed between embryonic and adult zebrafish SA node.

